# An integrated microfluidic platform for quantifying drug permeation across biomimetic vesicle membranes

**DOI:** 10.1101/523431

**Authors:** Michael Schaich, Jehangir Cama, Kareem Al Nahas, Diana Sobota, Kevin Jahnke, Siddharth Deshpande, Cees Dekker, Ulrich F. Keyser

## Abstract

The low membrane permeability of candidate drug molecules is a major challenge in drug development and insufficient permeability is one reason for the failure of antibiotic treatment against bacteria. Quantifying drug transport across specific pathways in living systems is challenging since one typically lacks knowledge of the exact lipidome and proteome of the individual cells under investigation. Here, we quantify drug permeability across biomimetic liposome membranes, with comprehensive control over membrane composition. We integrate the microfluidic octanol-assisted liposome assembly platform with an optofluidic transport assay to create a complete microfluidic total analysis system for quantifying drug permeability. Our system enables us to form liposomes with charged lipids mimicking the negative charge of bacterial membranes at physiological salt and pH levels, which proved difficult with previous liposome formation techniques. Furthermore, the microfluidic technique yields an order of magnitude more liposomes per experiment than previous assays. We demonstrate the feasibility of the assay by determining the permeability coefficient of norfloxacin across biomimetic liposomes.

## I. Introduction

Over the past decades, multidrug resistance (MDR) in microbial pathogens has developed into a serious threat for public health, leading to a global medical crisis^1^. In 2016, more than half (58.6%) of clinical *Escherichia coli* isolates in the European Union showed resistance to at least one of the antimicrobial groups under regular surveillance^2^. One of the major biochemical causes of antibiotic resistance is the reduced membrane permeability of drug molecules^3–6^. This problem is especially apparent for Gram-negative bacteria, as their cell envelope consists of a double membrane; drugs require seemingly contradictory chemical properties to overcome these two barriers to reach their cytoplasmic targets^4,6,7^. A deeper understanding of the mechanisms that govern passive drug transport across lipid membranes is therefore of great importance.

Bacterial cell membranes are very complex systems that are involved in numerous cellular processes^8^. Drug permeability studies have not only proven very difficult due to the small size of the bacteria, but also due to the convolution of active and passive effects that simultaneously take place in the membranes of living bacteria^6,9^. Many studies on membrane properties are therefore performed using lipid vesicles (or liposomes) as model systems^9^. Liposomes of several microns in size, referred to as giant unilamellar vesicles (GUVs), offer the advantages of having well-defined lipid compositions, being easy to image and also being more controlled systems for studying transport processes than a living bacterium. Due to these advantages, liposomes have been the subject of intensive research and their applications now reach far beyond the drug delivery^10^ and synthetic biology communities^11–13^.

Various methods to produce lipid vesicles have been developed^14^. Albeit offering good control over lipid composition, many technologies such as electroformation suffer from drawbacks such as low yield, batch-to-batch variability, low encapsulation efficiency and polydispersity; these problems are especially acute at high salt concentrations and when using charged lipids^15^. Microfluidic methods to form liposomes can overcome these limitations, but often require oil as the lipid carrying solvent, which must be removed in further procedures^16^. The recently developed microfluidic technique octanol-assisted liposome assembly (OLA) replaces the oil phase with the aliphatic alcohol 1-octanol^17^. A double emulsion of water in octanol self-assembles into a liposome with an octanol pocket attached to it upon production. The octanol pocket later pinches off within a few minutes, resulting in a liposome and a separated octanol droplet.

We have previously presented an optofluidic permeability assay which allows us to determine the permeability coefficient of electroformed GUVs to fluoroquinolone drugs in a direct, label-free manner^18^. The assay exploits the autofluorescence property of fluoroquinolones in the ultraviolet for label-free detection. GUVs are exposed to a drug solute in a controlled manner in a microfluidic device. We use ultraviolet video fluorescence microscopy to quantify drug uptake in the GUVs and report the permeability coefficient of the drug across the membrane composition of interest. We used this method to show the influence of lipid composition on fluoroquinolone transport^19^. Furthermore, we showed that the permeability of different drugs from the fluoroquinolone family can span over two orders of magnitude at different pH levels^20^. Finally, we used the assay to study the transport behaviour of proteoliposomes containing the *E. coli* outer membrane protein OmpF^21^, thus developing a direct, optical measurement for antibiotic flux through porins, which form an important route for drug molecules to translocate across the outer membrane of Gram-negative bacteria^5^.

Despite these advances, the technique suffered from the various drawbacks of off-chip liposome formation described above. In this paper, we present the successful integration of the OLA platform with our optofluidic transport assay. This lab-on-chip total analysis platform enables the continuous production and screening of liposomes on the same device. This enabled us to efficiently screen an order of magnitude more liposomes than in our earlier platform. Importantly, it also enables us to explore transport using physiological salt concentrations which was challenging using electroformed GUVs^22^. We also present an improved MATLAB analysis routine, which offers superior liposome detection, automatic channel recognition and more debugging options than our earlier platform. The method was validated by performing transport experiments of the fluoroquinolone norfloxacin through biomimetic PGPC liposomal membranes. We verified that the liposomes produced are indeed unilamellar by performing a dithionite bleaching assay.

## II. Experimental Design

### Microfluidic Chip Design

The microfluidic device integrates the channel features used for octanol-assisted liposome assembly (OLA) with a downstream T-junction geometry to form a complete lab-on-chip system for the continuous production and screening of liposomes. A schematic of the device is shown in Fig 1A.

**Figure 1.**
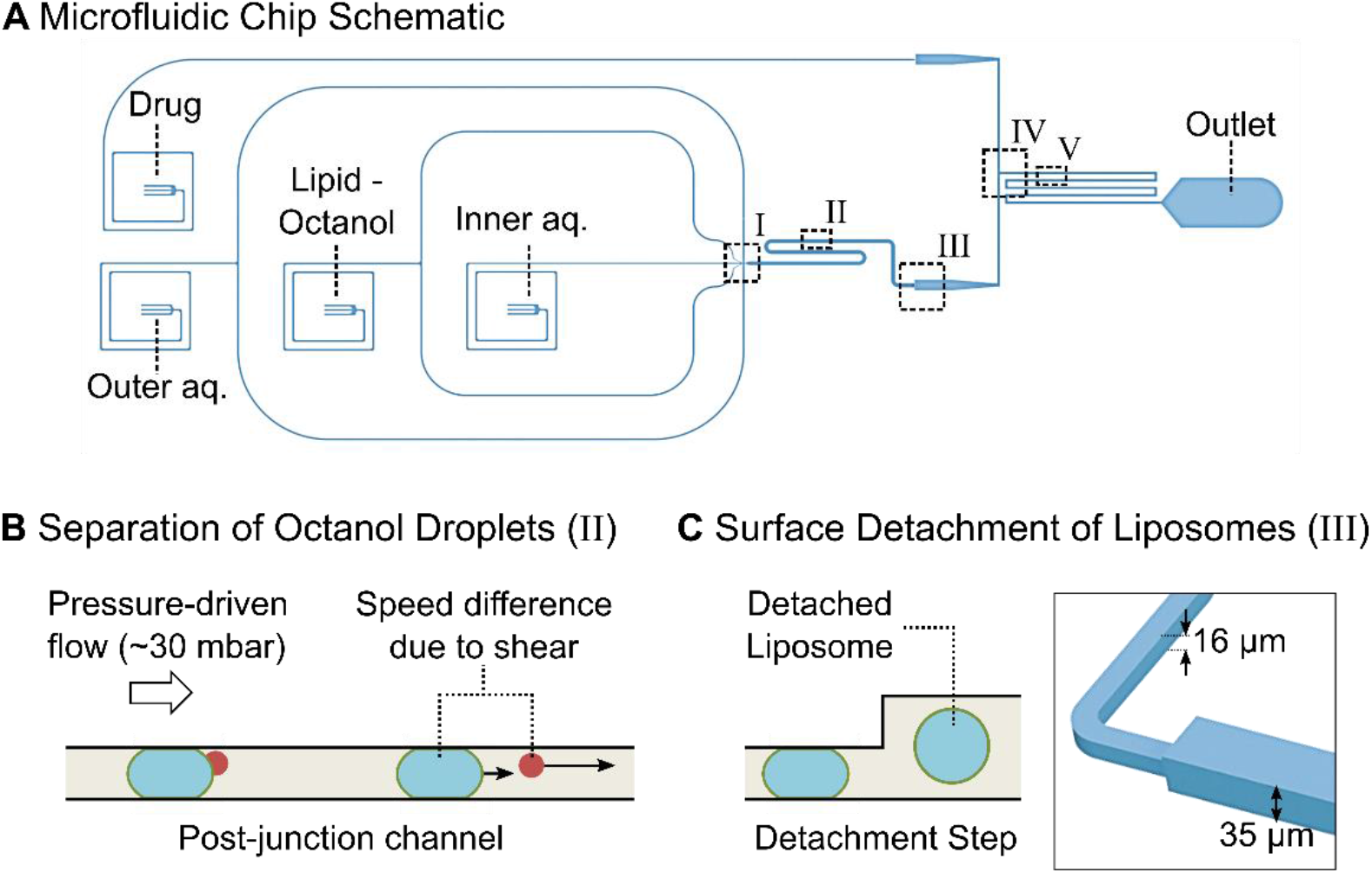
Microfluidic total analysis system for quantifying drug permeability across liposome membranes. **A**: The microfluidic chip features four inlets, one outlet and two different channel heights. The outer aqueous, inner aqueous and lipid-octanol inlets are needed for liposome production on chip. The fourth inlet is used to flush in the drug whose permeability is to be measured. The liposome production occurs at a 6-way junction, where the aqueous flows meet the lipid-octanol phase (I). The 1-octanol pocket which is initially attached to the liposome separates from it in the post-junction channel within minutes after production (II). After an increase in channel height (III), the liposomes are mixed with the drug solute (IV). The transport measurement takes place as the liposomes flow towards the outlet, immersed in a bath of the autofluorescing drug (V). **B:** Mechanism to separate the liposome population from the octanol droplets. The liposomes typically have radii of 15-18 μm upon production and octanol droplets of < 8 μm. The channel height post formation is lower than the diameter of the liposomes which leads to a significant difference in velocity for octanol droplets and liposomes as indicated by the arrows. The octanol droplets pass through the device first and are discarded at the outlet. **C:** Upon production, the liposomes’ diameters are larger than the height of the microfluidic channel. An increase in channel height from 16 μm to 35 μm frees the liposomes from the geometric confinement and enables transport measurements across the membrane without the risk of shear-induced leakage.

The 6-way junction where the liposomes are created is shown in Fig. 2A. The original design geometry^17^ was modified to fit the needs of the drug transport assay. The dimensions of the junction were scaled up by a factor of 2 to achieve liposome diameters of up to 35 μm. The larger channels furthermore lead to higher flow rates and higher liposome production rates. A side effect of the larger liposomes is that the budding off of the octanol pocket from the liposomes requires more time than for smaller vesicles. The budding off process has previously been reported to occur within a minute after production^23^, whereas this process is on the time scale of minutes using our chip geometry. The general mechanism of double-emulsion formation and separation into a liposome and a 1-octanol droplet remains the same as previously reported^17^.

**Figure 2.**
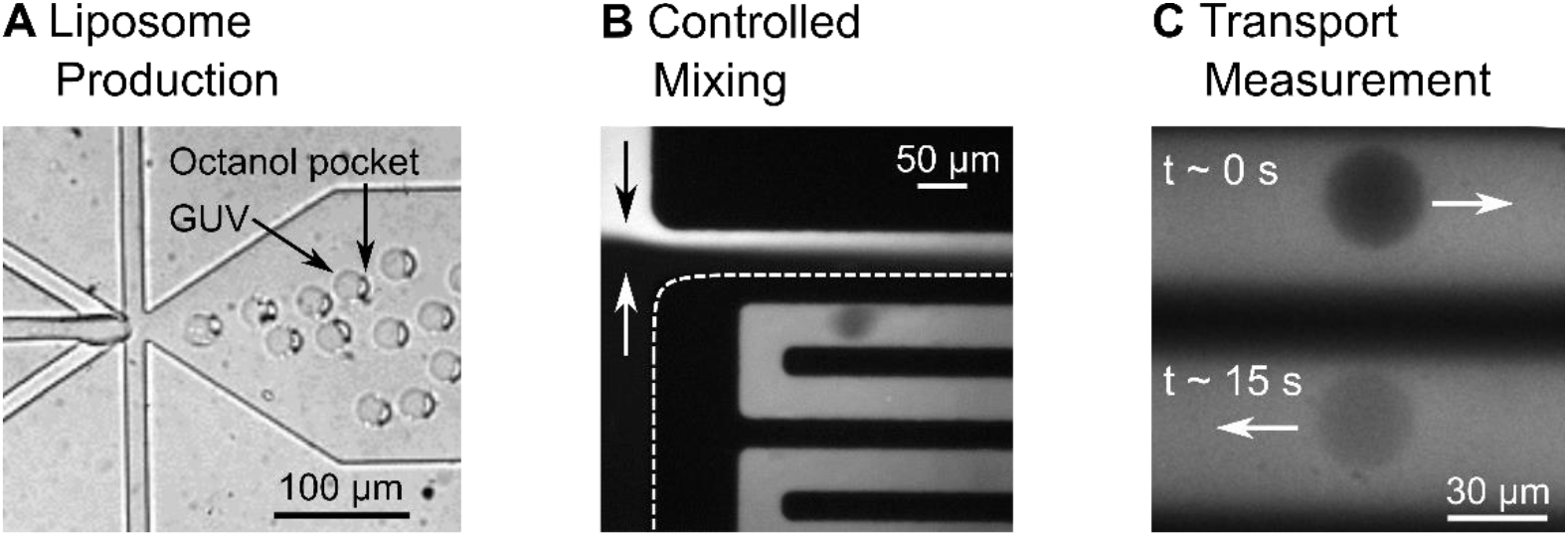
Liposomes at different positions in the microfluidic chip. **A:** Liposome assembly at the 6-way formation junction. A 1-octanol pocket is initially attached to the liposomes. The liposome and the octanol pocket separate further downstream in the post-junction channel. **B:** The liposomes experience a spontaneous exposure to a drug solute at a T-junction where the two flows mix in a controlled manner. **C:** The liposomes, surrounded by the autofìuorescing drug (λ_ex_ = 350 nm), can be monitored at different parts of the channel, corresponding to different times that the liposome has been exposed to the drug. The increase in liposome intensity as the fluorescing drug diffuses across the membrane is used to determine the permeability coefficient of the drug across the lipid membrane under investigation.

The generated liposomes have diameters larger than the height of the post-junction channel. These liposomes therefore initially shear along the PDMS walls, as they flow downstream. To ensure that the shear does not compromise the membranes during the subsequent transport measurements, an increment in channel height was introduced, increasing the channel height from 16 μm to 35 μm (Fig. 1C). Post the step, the liposomes flow towards the mixing junction without shearing.

Reaching the T-junction, the liposomes are exposed to a drug solute as they flow towards the outlet channel (Fig. 2B). Drug transport across the liposome membrane is tracked label-free by exploiting the autofluorescence of the drug in the UV region (λ_ex_ = 350 nm). The liposomes initially appear dark on a bright background due to the lack of fluorescent drug molecules within them. As the fluorescent drug molecules diffuse through the membrane, the liposomes get brighter, as seen in Fig. 2C. We record and analyse the increase in fluorescence intensity using a previously established analytical protocol to quantify the permeability coefficient of the drug molecule for the specific lipid composition under investigation^18,20^.

### Separation of Octanol Droplets and Liposomes Based on Flow Speed

The shearing of the liposomes in the post-junction channel is exploited to separate the liposome population from the octanol droplets. This separation mechanism, visualized in Fig. 1B, is based on the difference in flow speeds between the liposomes and the droplets. Upon production, the liposomes typically have radii of 15-18 μm, whereas the octanol droplets have radii of under 8 μm. The channel height of 16 μm before the step causes deformation and shearing of the liposomes with the PDMS and slows their flow speed down considerably. In comparison, the smaller octanol droplets do not shear and therefore possess a higher velocity. Liposomes typically move with speeds of ~0.05 mm/s, whereas the droplets move at ~0.2 mm/s in this region of the chip. By stopping the lipid-octanol flow after a desirable number of liposomes is created and filling the post junction channel, this mechanism leads to a separation of the two populations. The octanol droplets get flushed through, while the GUVs remain in the chip. After reaching the step, the liposomes are no longer pressed against the channel wall, reconfigure to an expected isotropic spherical geometry and encounter the drug flow at the T-junction.

## III. Materials and Methods

### Chip Fabrication

The microfluidic chips are made of polydimethylsiloxane (PDMS) using established photo- and soft-lithography techniques. The master mold is generated by spin coating a thin layer of SU-8 2025 photoresist (Chestech, UK) on a 4-inch Silicon wafer (University Wafer, USA). The wafer is pre-baked on a hot plate at 65°C for 1 min and at 95°C for 6 min. The structures are imprinted on the substrate using a table-top laser direct imaging (LDI) system (LPKF ProtoLaser LDI, Germany). The LDI system exposes the structures specified in the software directly with UV light, causing the photoresist to crosslink and solidify. After exposure, the wafer is post-baked for 1 min at 65°C and for 6 min at 95°C. The substrate is developed by rinsing the wafer with propylene glycol monomethyl ether acetate (PGMEA), which removes the unexposed photoresist and leaves the desired UV-exposed structures on the substrate. The wafer is then hard baked for 15 min at 120°C. The multi-height feature is achieved by performing this photo-lithography process twice on the same silicon wafer with different layers of photoresist with varying heights. Feature heights of 16 μm and 35 μm respectively were obtained by spinning the photoresist at 3800 rpm and 1800 rpm respectively (WS-650-23NPP, Laurell Technologies, USA) for 60 s with a ramp of 100 rpm/s. The anchoring tool of the direct laser writer is used for aligning the features in the two designs.

The PDMS microfluidic devices are made by using the silicon master as a mold. Liquid PDMS (Sylgard 184, Dow Corning) is mixed in a 9:1 ratio with the curing agent and desiccated to remove air bubbles. It is then cast into the mold and cured for 60 min at 60°C. Fluid access ports of 0.75 mm diameter for the inlets and 1.5 mm diameter for the outlet are punched into the chip using biopsy punches (WPI, UK). The PDMS chip is then plasma-bonded to PDMS-coated cover slips using a standard plasma bonding protocol (100 W, 10 s exposure, 25 sccm, plasma oven from Diener Electric, Germany).

The surfaces of the outlet channel are rendered hydrophilic by flushing the channel with a polyvinyl alcohol (PVA) solution for 15 min (50 mg/mL, 87-90% hydrolysed molecular weight 30,000-70,000 Da, Sigma-Aldrich) via the outer aqueous inlet, using a previously established protocol^17^. Post treatment, the PVA is removed from the channels by applying suction with a vacuum pump (Gardner Denver Thomas GmbH, Germany) after which the microfluidic device is baked in the oven at 120°C for 15 min.

### Optical Setup

A custom-built UV epifluorescence setup is used to induce autofluorescence in the drug molecules and to capture the experimental video data. White light from a broadband light source (EQ99FC, Energetiq, USA) enters a monochromator (Monoscan 2000, OceanOptics, USA) where the excitation wavelength (λex = 350 nm) for the target drug molecule is selected. The UV light passes through a Köhler illumination pathway and illuminates the microfluidic device via a quad band dichroic mirror (BrightLine full-multiband filter set, Semrock, USA) and a microscope objective. The fluorescence signal is detected by an EMCCD camera (Evolve 512 Delta, Photometrics). The camera and recording settings (exposure 10 ms, bin 2, gain 150) are controlled using the open source software μManager 1.4 ^24^. A 60× water immersion objective (UPLSAPO NA 1.2, Olympus) is used for data recording, whereas lower magnifications (4×, 10×, 20×, Plan Achromat, Olympus) are used for optimising the vesicle formation and PVA treatment of the chip.

### Solution Compositions and Flow Control

The base solutions for the OLA aqueous phases consist of 200 mM sucrose and 15% v/v glycerol in buffer. In accordance to previously published protocols, the outer aqueous phase additionally contains 50 mg/mL poloxamer Kolliphor P-188, which facilitates the initial double-emulsion formation^23^. Experiments are performed in two different chemical environments. Transport is measured in phosphate-buffered saline (pH 7.4) which mimics physiological salt and pH conditions. Additionally, experiments using a 5 mM acetic acid buffer (pH 5) are performed as controls for membrane stability. At pH 5, norfloxacin molecules are primarily in their positively charged form and hence show low permeability through lipid bilayers^18,21^. The drug solute is prepared by diluting a 48.5 mM norfloxacin stock solution to a final concentration of 2 mM with the aqueous base stock. All pH levels are adjusted and checked using a digital pH meter (Hanna Instruments, UK). The lipid-octanol phase is obtained by dissolving a 100 mg/mL lipid stock mixture with 1-octanol to reach a final concentration of 2 mg/mL. The lipid stock is a 3:1 mixture of DOPC (1,2-dioleoyl-sn-glycero-3-phosphocholine) and DOPG (1,2-Dioleoyl-sn-glycero-3-phosphorac-(1-glycerol) sodium salt) in 100% Ethanol. We used the 3:1 DOPC:DOPG composition to mimic the anionic charge of bacterial membranes^25,26^. All chemicals are obtained from Sigma-Aldrich, unless stated otherwise.

The liquid flows in the microfluidic device are controlled with a pressure-driven microfluidic pump (MFCS-EZ, Fluigent). The fluids are stored in Fluiwell-4C reservoirs (Micrewtube 0.5mL, Simport) and enter the microfluidic chip via a polymer tubing (Tygon microbore tubing, 0.020’’ x 0.060’’ OD, Cole Parmer). Cut dispensing tips (Gauge 23 blunt end, Intertronics) are used as metal connectors between the tubing and the chip.

### Experimental Protocol

The microfluidic assay involves a 2-stage protocol. The first stage involves adjusting the pressure-driven flows of the liquids to obtain stable liposome production, as reported previously^17^. Typically, pressures of ~40 mbar for the inner aqueous, lipid-octanol and drug inlet phases, and ~70 mbar for the outer aqueous phases lead to a stable production of liposomes. After the post-formation channel has been filled with 150-200 liposomes, the aqueous flows are reduced (IA and OA input pressures at this stage are typically reduced to around 15 mbar each), and the lipid-octanol flow is stopped completely. Due to the difference in flow speeds described in Fig. 1B, the octanol droplets flow out of the chip leaving a population of octanol free liposomes in the post-formation channel. Throughout the entire process the T-junction is regularly monitored to ensure that the drug and OLA flows mix equally and that no octanol enters the drug channel.

The next stage involves the drug permeability measurement, using the same measurement principles previously employed in our laboratory^18–21^. The field-of-view is changed to the channel network post the T-junction, shortly after the liposomes encounter the drug flow (Fig. 2C). Using a 60× water immersion objective (UPLSAPO, NA 1.2, Olympus), two sections of the channel are monitored simultaneously with sufficient resolution to study the transport process. Since the liposome is dispensed in the drug solution flowing along the channel, the different positions in the channel correspond to different drug exposure times of the liposome. The IA and OA pressures are maintained at 15 mbar each, resulting in flow speeds in the channel of about 0.5 mm/s. One measurement is taken just after the drug and the vesicle flows meet (t ≈ 0 s), the other one typically after the liposomes have been exposed to the drug for ~15 seconds. Measurements of up to 40 seconds of drug exposure can be obtained at these flow rates by recording the liposomes at the end of the channel network, just before the outlet reservoir. This can be further increased by increasing the length of the mixing channel^18^.

After all the liposomes have been flushed through, the liposome formation can be restarted, and the experiment repeated. The number of repeats is limited by the quality of the PVA coating and/or accumulation of lipid and octanol aggregates in the chip. We achieved up to four such repeats on a single microfluidic device.

### Data Processing and Permeability Calculation

The data is analysed using our previously established permeability model^18,20,21^. The videos obtained are processed using the MATLAB script provided in the Supplementary Materials. Similar to our previously reported analysis routine, the script extracts the radius, the speed, the circularity and the intensity values of the liposomes passing through the channels. We have updated the image analysis routine, which now additionally features superior liposome detection, automatic channel recognition and debugging options that are explained in detail in the Supplementary Information.

The permeability coefficient is obtained by the equation^18^:

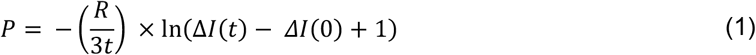

Where the variable *R* is the radius of the liposome, *t* is the time of its exposure to the antibiotic and *ΔI* is the normalized autofluorescence intensity difference between the interior (*I_in_*) and the exterior (*I_out_*) of the liposome:

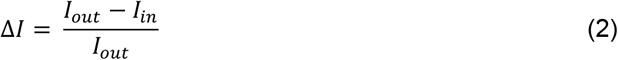

The difference in the normalized intensity of the liposomes at different drug exposure times can be used as a direct readout of drug flux into the liposomes^18^. Due to the initial lack of fluorescing drug molecules inside the liposome, *ΔI*(*t* = 0) has a high value, correlating to a large difference in fluorescence intensity between the liposome (*I_in_*) and the background (*I_out_*). In contrast, Δ*I*(*t*) at the later time point has a lower value consistent to a lower difference in fluorescence intensity between the liposome and the background, due to the influx of fluorescing drug molecules. Further explanations and a derivation of Equation (1) have been published in our previous work^18,20^.

## IV. Test for Membrane Unilamellarity

Liposomes produced by OLA have previously been tested for their unilamellarity via the incorporation of the pore forming toxin α-hemolysin^17^. Additionally, the antimicrobial peptide cecropin B was found to permeabilise and lyse OLA-produced liposomes, again suggesting that the liposomes are unilamellar^27^. As an additional, quantitative technique to verify the unilamellarity of the OLA-produced liposomes, we extracted the liposomes from our microfluidic device and subjected them to a dithionite bleaching assay. If brought in contact with it, the membrane-impermeable anion dithionite reduces and thereby irreversibly bleaches nitrobenzoxadiazole (NBD). By subjecting unilamellar liposomes containing NBD-labelled lipids to a dithionite solution, the fluorescence intensity of the liposome drops to half of the initial value, due to the bleaching of the outer leaflet of the bilayer membrane^28,29^.

The composition of the inner and outer aqueous solutions used for liposome production in this experiment is identical to the solutions of the drug transport assay described above. The lipid-octanol phase additionally contains 0.01 mg/mL of a fluorescently labelled NBD-PC lipid (1-palmitoyl-2-{6-[(7-nitro-2-1,3-benzoxadiazol-4-yl) amino] hexanoyl}-sn-glycero-3-phosphocholine, Avanti Polar Lipids, USA). A standard PDMS chip with the design shown in Fig. 1 is used to extract the liposomes off chip. However, instead of applying a Ø 1.5 mm punch at the outlet, a Ø 4 mm hole is punched at the detachment step in this case. The Ø 4 mm outlet serves as a fluid reservoir where the liposomes are collected. After the pressures of the microfluidic pump have been adjusted to achieve a stable liposome production, an additional 20 μl of the IA solution is pipetted into the reservoir, which aids the separation of liposomes and octanol droplets in the reservoir. Due to the lower density of octanol, the droplets rise to the surface of the reservoir. A larger fluid volume furthermore facilitates liposome extraction. After 1 hour of liposome production, 15 μl of the liposome suspension is extracted from the reservoir using a wide bore pipette tip.

The liposome suspension is added to a microscopy chamber (Grace Bio-Labs FlexWell^™^, Sigma Aldrich) on a BSA-coated coverslip containing 35 μl of a solution containing PBS, 200 mM glucose and 15% v/v glycerol. The liposomes are left for 1 hour to sink and settle at the bottom of the chamber which facilitates imaging. A dithionite solution stock (1M sodium dithionite in Tris pH 10 buffer, Sigma Aldrich) is diluted in the glucose buffer to a final dithionite concentration of 15 mM. 30 μl of the solution is added to the liposome suspension after the imaging is started^29^. Imaging is performed on a confocal microscope (Olympus IX83, FV10-MCPSU laser system, 20x objective UPLSAPO Olympus, 5 s frame interval). Image analysis is performed using the open source software Image J.

The intensity traces of the liposomes normalized to their initial intensity value are depicted in Figure 3. Fig. 3A shows the intensity drop upon addition of dithionite, whereas Fig. 3B shows the results of the bleaching control experiment upon addition of buffer without dithionite. The mean intensity of the observed liposomes is shown as black line with the standard deviations depicted in grey. The intensity of the liposomes subjected to dithionite (N = 10) drops to half of the initial intensity after about 500 seconds and then stays steady at that value. These results suggest that the liposomes indeed consist of a single lipid bilayer, whose outer leaflet is bleached by the dithionite^29^. Since the dithionite anion cannot penetrate the membrane, the inner leaflet of the membrane is not affected by the dithionite and remains fluorescent. The control experiment without dithionite in Fig 3B shows a stable intensity signal over the timespan of the entire experiment (N = 3). This suggests that photo bleaching does not play a significant role in the observed drop in the intensity and that we are indeed observing the bleaching of the outer leaflet of a single bilayer.

**Figure 3.**
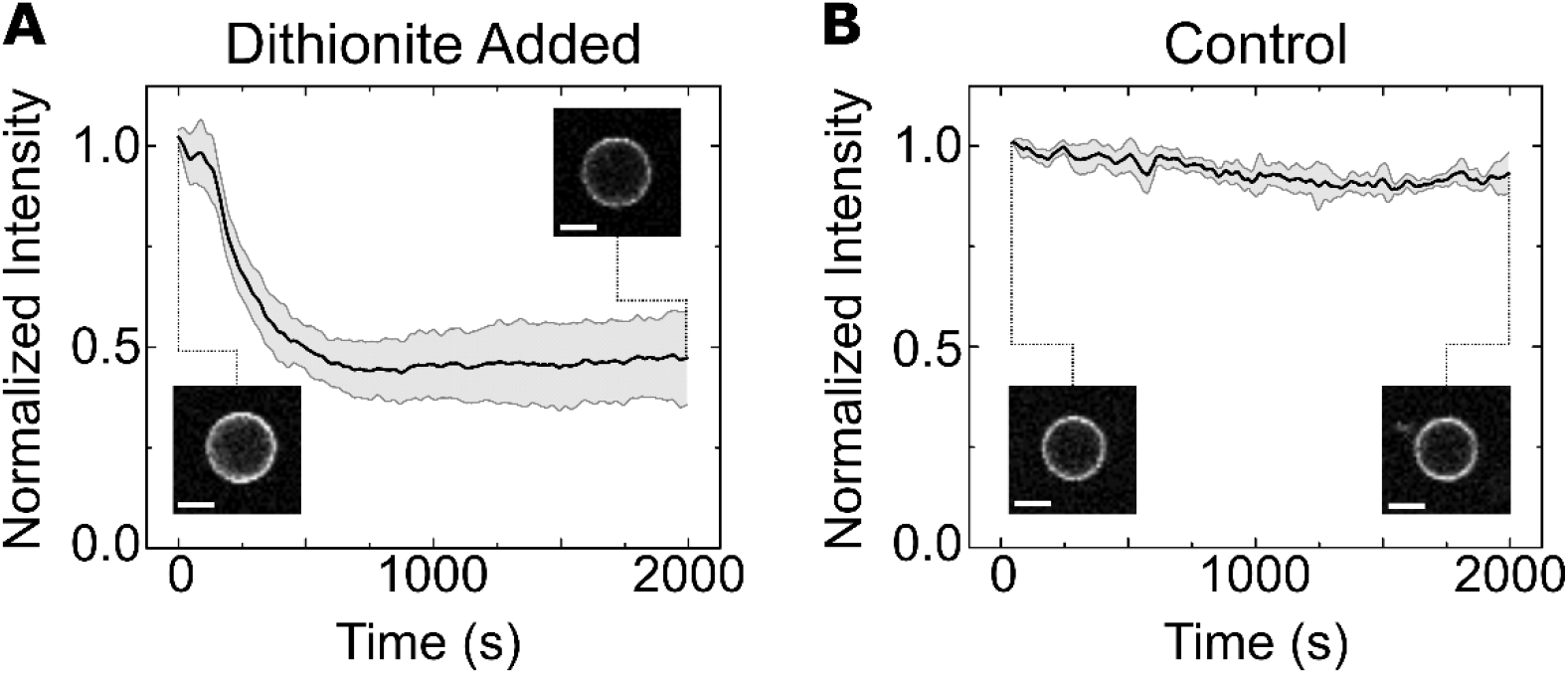
Normalized intensity traces of the liposome membranes. **A:** Intensity drop of the liposome membranes (N=10) upon addition of dithionite. The mean intensity of the observed liposomes is shown in black with the standard deviations depicted in grey. The drop to half of the initial value is caused by the bleaching of the outer membrane leaflet by the dithionite. **B:** Liposome intensity (N = 3) stays stable throughout the entire experiment upon addition of buffer without dithionite, suggesting a negligible effect of photo bleaching. The insets show a representative liposome at the beginning of the measurement and after 2000 s. The scale bar shows 10 μm.

## V. Results and Discussion

### Passive Diffusion Measurement of Norfloxacin

Drug transport measurements were performed in two different chemical environments. One set of measurements was performed using physiological salt and pH conditions (PBS). The second set was taken as a control in an acetic acid buffer at pH 5. The majority of norfloxacin molecules are positively charged at pH 5 whereas the proportion of uncharged molecules is increased at pH 7.4^18,20,21,30^. The liposomes consisted of PCPG lipids in a 3:1 ratio, a mixture which is commonly used as a model for bacterial membranes due to its negative charge^25,26^.

The scatter plots in Fig. 4 show representative results from such experiments. The black data points mark the normalized intensity difference levels *ΔI*(0) of individual liposomes at the first measurement point, when the liposomes have just encountered the drug flow and hence do not contain any drug molecules within them. The coloured data points mark the normalized intensity difference *ΔI*(*t*) of the individual liposomes after they have been exposed to the drug for the time indicated inset. The liposomes in PBS show a substantial drop in *ΔI* after being exposed to the drug, whereas the measurements at the different time points overlap when the studies are conducted at pH 5.

**Figure 4.**
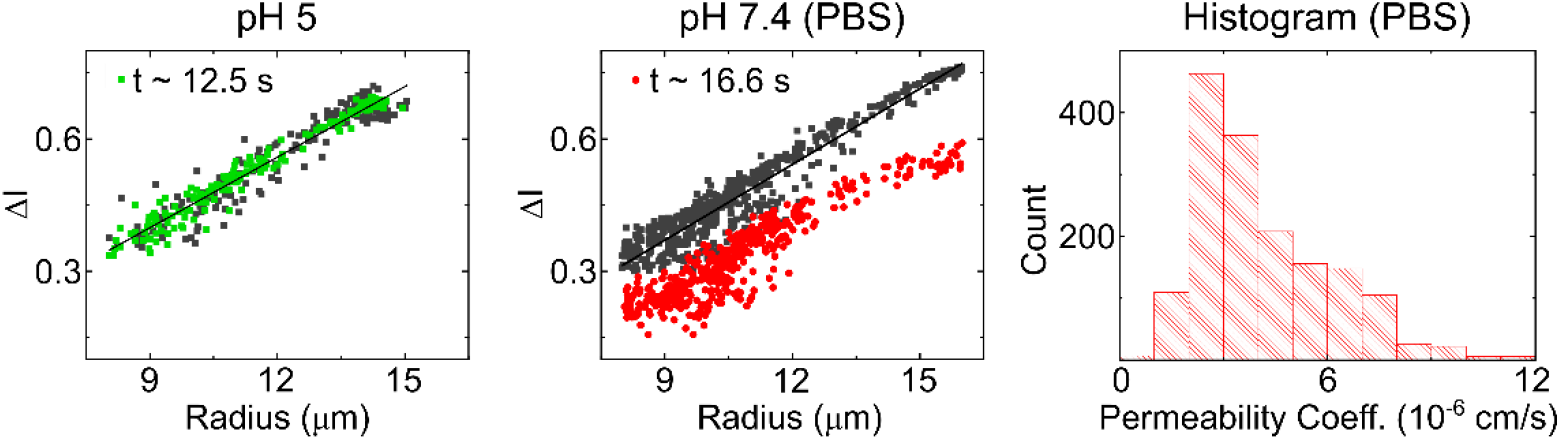
Norfloxacin diffusion experiments across PCPG liposomal membranes. The scatterplots show representative drug uptake experiments at pH 5 and pH 7.4, respectively. Each point in the scatter plots corresponds to the measured normalized intensity difference of a liposome. The dark points are obtained at t ≈ 0, just after the liposomes encounter the drug flow. The coloured points show the value at the second measurement position, after exposure to the drug for the time indicated in the panel. The gap between the black and red points for the measurement in PBS indicates norfloxacin membrane permeability. The histogram shows the distribution of all permeability measurements combined at pH 7.4 (N = 1620, 5 technical repeats). The distribution has a mean value of 4.13 ± 0.05 x 10^−6^cm/s (mean ± std. error of mean) and a median value of 3.57 x 10^−6^ cm/s.

Since the drop in *ΔI* is a direct result of the influx of fluorescent molecules, we can conclude that no significant transport occurs at pH 5 over the observed timespans, whereas there is substantial norfloxacin transport at physiological pH and salt conditions. The technical repeats of this experiment shown in Fig. S3 and Fig. S4 (ESI) report similar results. Expanding the exposure and observation time for the pH 5 measurement to 40 seconds yields the same result of no significant transport, in line with our previous observations. This also confirms that the liposome membranes are not being compromised due to shear or any other interactions in our device.

The observed transport behaviour is to be expected, as the norfloxacin molecule is predominantly uncharged or zwitterionic at neutral pH, whereas the molecule is in a charged state at pH 5 ^31^. Lipid membranes are generally regarded as largely impermeable to ions and highly charged molecules, since these cannot cross the hydrocarbon section of the lipid bilayer easily. In nature, these transport processes are governed by transmembrane proteins such as porins or ion channels^32^.

The spread in vesicle radius observed in Fig. 4 is a result of the method used to separate the liposomes from the octanol droplet population. As illustrated in Fig. 1B, the mechanism is based on a difference in flow velocity in the channel due to shearing of the liposomes. Throughout their travel to the step in channel height, the membranes are susceptible to small disruptions due to shear. The disruptions can lead to shrinking and separation of the liposomes. This spread in liposome size is illustrated in Fig. S2 (ESI). The gradient of the curve that the scatter points lie on is a result of the fact that our optical measurement is not a confocal measurement. The fluorescent drug solute surrounding smaller liposomes therefore leads to a lower apparent *ΔI,* compared to larger liposomes (for a detailed explanation, please refer to Cama *et al*.^18^). The analysis protocols are hence identical to our previously established technique.

The resulting permeability coefficients of all the experiments performed in PBS are combined in the histogram in Fig. 4. The total number of liposomes in the histogram is 1620 and results in an overall average permeability coefficient of 4.13 ± 0.05 x 10^−6^ cm/s (mean ± std. error of mean) and a median value of 3.57 x 10^−6^ cm/s. The values were obtained from 12 experiments in 5 technical repeats. For the pH 5 measurement, five experiments on two technical repeats were performed. An experiment is defined as a continuously acquired dataset measured from one batch of liposomes produced on the microfluidic device. As described above, up to four experiments were performed on one microfluidic device, by restarting the liposome production after completion of one measurement. Each set of experiments performed on an individual device is defined as a technical repeat.

The value obtained from the transport measurements matches the values previously obtained in our group using the established electroformation liposome production technique. Purushothaman *et al.* measured the permeability coefficient of norfloxacin through PGPC liposomes (30:70 ratio) in a 5 mM phosphate buffer solution at pH 7.0 to be 4.3 ± 0.2 x 10^−6^ cm/s^19^. The main advantage of the on-chip OLA technique over electroformation is the high-throughput liposome production and its compatibility with physiological salt concentrations. The liposome production efficiency allows us to perform tests on 1620 liposomes here compared to the 138 liposomes in the experiments using electroformation. Furthermore, the possibility of producing liposomes at physiological salt concentrations allows us to mimic the natural environment of a bacterial cell more closely. Any possible remaining traces of solvent^23^ (1-octanol, poloxamer P-188) do not seem to alter the transport properties of the membrane, as seen by the similarity to the previously acquired result.

## VI. Conclusion

In this paper, we presented an integrated microfluidic platform for quantifying drug permeation across biomimetic membranes. We combined an on-chip liposome formation technique (OLA) with a downstream T-junction for the controlled exposure of liposomes to a drug solute. Norfloxacin transport through biomimetic PGPC (1:3 ratio) liposomes was measured at physiological salt and pH conditions and yielded a permeability coefficient of 4.13 ± 0.05 x 10^−6^ cm/s (mean ± std. error of mean) and a median value of 3.57 x 10^−6^ cm/s, which is in good agreement with previous measurements^19^.

Since our method directly quantifies the permeability coefficient of the drug across the specific membrane of interest, our technique offers an alternative to traditional drug transport assays such as octanol-partition coefficients, or the parallel artificial membrane permeability assay (PAMPA) which suffer from multiple drawbacks^33,34^. Our method also profits from recent advances in the development of fluorescent antibiotics, as these provide another potential tool for studying and understanding drug transport processes that will lead to further insight into drug-membrane interactions^35^. Importantly, thanks to the microfluidic character of our method, we require only very small reagent volumes in the microliter range for our measurements^36^.

Another advantage of the integrated on-chip technique presented here over our previously published optofluidic permeability assay lies in the benefits of controlling liposome formation with OLA. OLA allows the formation of large numbers of liposomes with physiological salt concentrations and with complex lipid mixtures. Other techniques such as electroformation suffer from very low yields in this environment^15^, or in the case of other microfluidic techniques, require extensive procedures to remove oil remnants associated with the production^17^. Moreover, OLA allows for the efficient encapsulation of desired solutes inside the vesicles upon production^17,27^. This makes it an interesting technique for biosensor-based approaches to detect drug molecules^37^.

The microfluidic platform can be expanded to study more complex membrane compositions and proteins. The technique could therefore be used for the formation and study of proteoliposomes, potentially offering an alternative to current reconstitution techniques^38^. Membrane transporters are an interesting field of study on multiple levels. On one side, they play a crucial role in drug uptake in bacteria and mutations of membrane proteins have been associated with antibiotic resistance^21,39^. On the other side, membrane transporters in the liver and kidney control the systemic clearance of drugs^40^. They therefore govern drug concentration profiles in the body with major effects on drug efficacy. Both these and other membrane transport-related questions can be addressed with our platform.

## VII. Acknowledgements

MS is funded by the Friedrich-Naumann-Foundation for Freedom, JC acknowledges funding from the BBSRC. KAN acknowledges support from an EPSRC CASE award with the National Physical Laboratory, Winton Programme for the Physics of Sustainability, Trinity-Henry Barlow Scholarship and the ERC. DS is funded by the Winton Programme for the Physics of Sustainability and the EPSRC. KJ was supported by the Erasmus Plus student exchange programme. SD and CD acknowledge support from the ERC Advanced grants SynDiv (no. 669598) and the Netherlands Organisation for Scientific Research (NWO/OCW), as part of the Frontiers of Nanoscience program. UFK acknowledges support from an ERC consolidator grant (Designer-Pores 647144).

