## Supplementary Information for "An integrated microfluidic platform for quantifying drug permeation across biomimetic vesicle membranes"

### 1. Experiment Dataset

The scatter plots of the experiments performed in PBS at pH 7.4 are provided in Fig. S3a and S3b below. The data for the experiments at pH 5 is provided in Fig. S4. The average time between the measurement positions is indicated in the inset together with the number of liposomes that have been detected at the second measurement point.

It should be noted that when calculating the permeability coefficient using Equation (1), the permeability was not obtained by comparing a unique vesicle at two different time points. The value of $\Delta I(0)$ was recalculated using a linear function that was fitted to the data points as a function of the radius. This fitted function is featured as a linear line in the scatter plots. The function is used to recalculate $\Delta I(0)$ for the vesicle radius detected at the second time point. This method still allows for the individual calculation of the permeability coefficient for each vesicle but abolished the need for identifying the same vesicle in multiple parts of the channel. Thus, the field of view can be shifted to any position in the channel network correlating with the desired time of drug exposure. The liposome intensities measured at this position can then be compared to the previously acquired intensities of the liposomes at the initial measurement position.

### 2. MATLAB Analysis and Data Filtering

The MATLAB routine used for the analysis is a modification of our previously presented method. For a detailed description of the procedure, please refer to the annotated code in the supplementary materials and our previous publications[1,2]. The following section will present the improvements compared to our old code.

A liposome entering the field of view is detected by observing a drop in average intensity of the frame. The detection sensitivity could be improved compared to our old method by screening the individual channels for an intensity drop, rather than screening the entire frame. The large number of liposomes generated by OLA furthermore makes it necessary to account for the case that several vesicles are visible in the channel at the same time. This case rarely occurred previously, due to the lower frequency of vesicle events. The updated version therefore includes a liposome identification element that ensures that the extracted vesicle information is not compromised if multiple vesicles are visible in the field of view at the same time. Additionally, the debugging options for the data were improved. The script now exports an image of the automatically identified channels, a plot of the channel intensity used to identify a liposome entering the field of view, and a debugging overview of the identified vesicle events shown in Fig. S1.

* Cavendish Laboratory, University of Cambridge, JJ Thomson Avenue, Cambridge CB30HE, United Kingdom
† Department of Biophysical Chemistry, University of Heidelberg, Im Neuenheimer Feld 253, D 69120, Heidelberg, Germany
‡ Max Planck Institute for Medical Research, Department of Cellular Biophysics, Jahnstraße 29, D 69120, Heidelberg, Germany
§ Department of Bionanoscience, Kavli Institute of Nanoscience, Delft University of Technology, Van der Maasweg 9, 2629 HZ Delft, The Netherlands

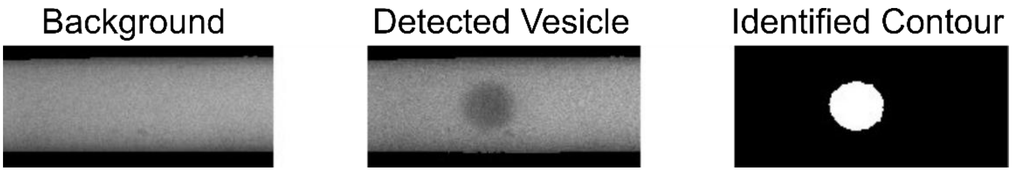


***Figure S1. Overview allowing for quick debugging of vesicle events.** The background image used for the normalization of $\Delta I(t)$ can be screened for correct identification as well as possible contamination with lipid aggregates or the presence of liposomes, which distort the background subtraction. The detected vesicle can be examined for its integrity and the complete detachment of the octanol pocket. Furthermore, the correct identification of the liposome contour can be checked.*

The debugging overview shows three images which together give a good indication of the validity of the liposome. Due to an ID stamp in the picture name, every single vesicle event can therefore be easily screened for errors. The first image shows an image of the background, constructed by averaging the images of the frames before and after the liposome event. It is important to avoid errors in this image, as the background intensity values used to calculate the normalized intensity $\Delta I$ are taken from this image. Incorrect background identification or impurities in the frame, such as lipid aggregates or full liposomes, can alter the normalization. The middle image shows the grayscale image of the detected vesicle. It can be screened for its integrity, contact/shear with the channel wall, or for the presence of an octanol pocket. Finally, the third image shows a binary image of the identified vesicle contour. It allows for easy debugging if the shape of the liposome was identified incorrectly. This is important for the correct measurement of vesicle radius and intensity.

The dataset was furthermore filtered to exclude any remaining octanol droplets, lipid aggregates, out of focus vesicles and false positives, as these skew the measurement. For this purpose, the data set of every scatter plot was filtered to only include liposomes of a radius 8 µm < $R$ < 16 µm, vesicles with initial $\Delta I(0)$ of > 0.3 and with a velocity < 1.0 mm/s. Furthermore, liposomes which still contained an octanol pocket attached to them or which showed an error in the background subtraction were also filtered out. This was performed using the improved debugging feature explained above.

### 3. Spread in Liposome Radius

One of the features of OLA is the monodispersity of the liposomes upon production. The spread in liposome size observed in these experiments is a result of the shear subjected to the liposomes, as they flow along the channel. The spread is visualized in Fig. S2. It shows liposomes at different distances from the formation junction, indicated in the figure. The histograms show a clear increase in polydispersity the further away the liposomes are from the formation junction and thereby the longer they are subjected to shearing.

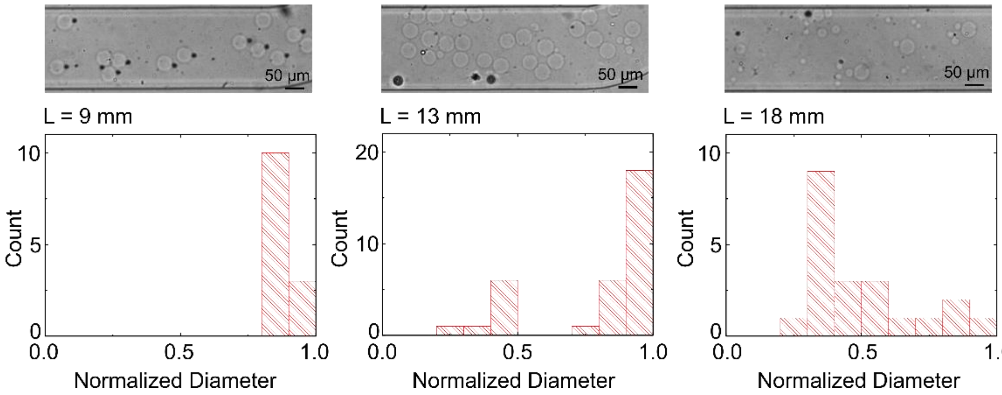


***Figure S2. Size distribution of liposomes in different parts of the channel.** The bright field images in the top row show liposomes at the indicated distances from the formation junction. The histograms show an increase in size spread the further away the liposomes are from the junction and thereby the longer they are subject to shearing with the PDMS chip.*

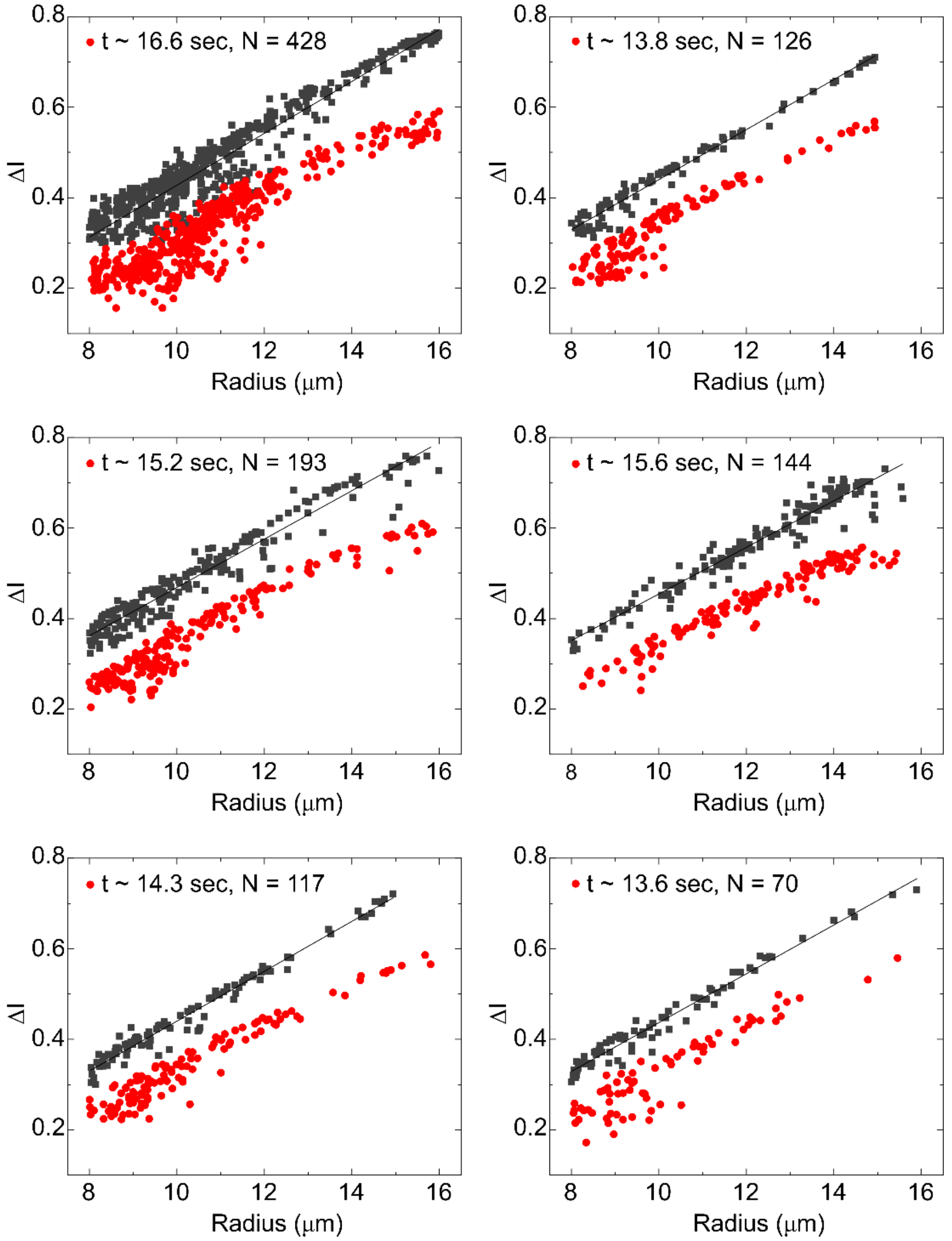


***Figure S3a. Scatter plots of ΔI vs R for norfloxacin in PBS with 200 mM sucrose at pH 7.4.*** *Significant transport of the drug through the PGPC vesicles can be detected in all experiments, visible by the gap in ΔI between the two time points.*

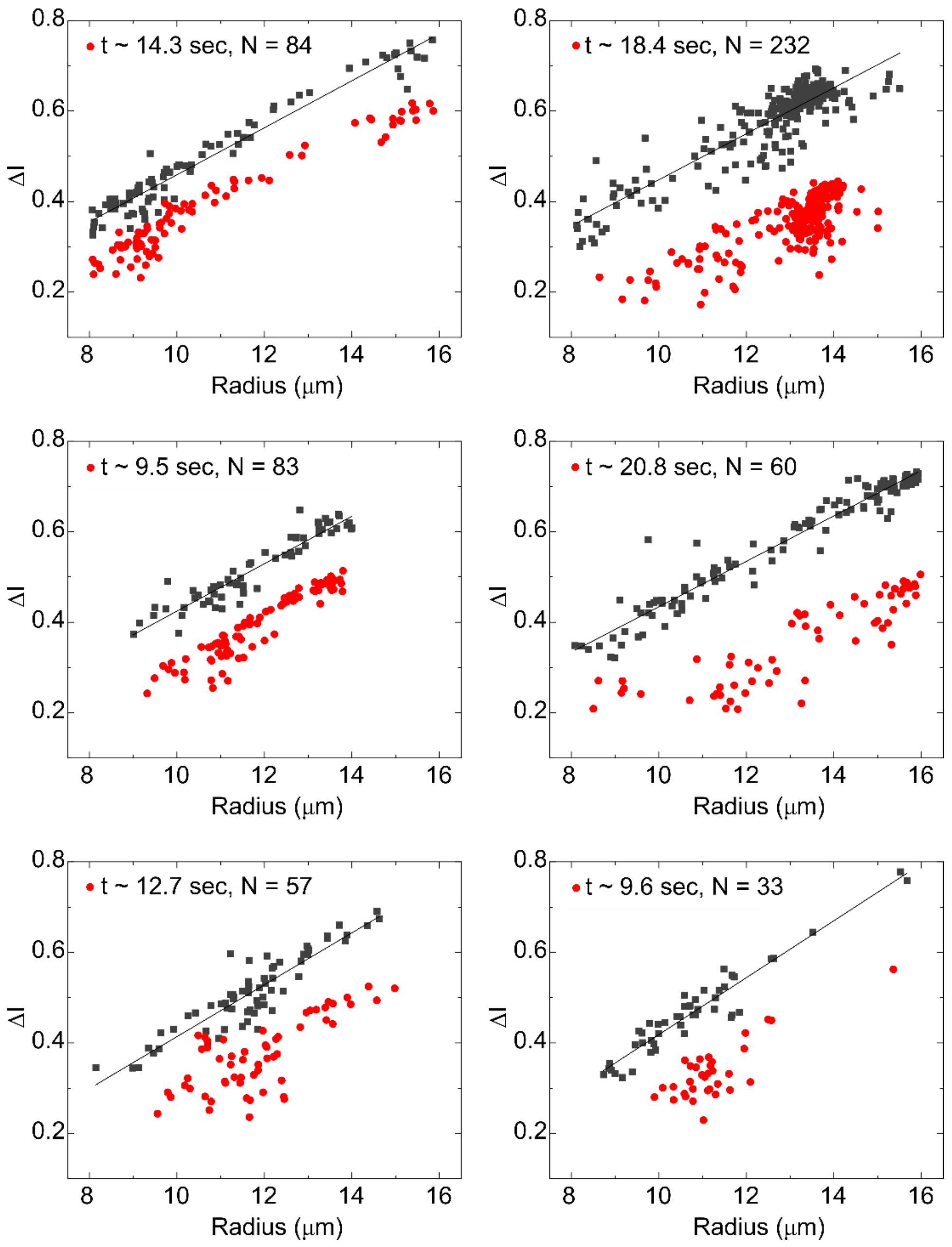


***Figure S3b. Scatter plots of ΔI vs R for norfloxacin in PBS with 200 mM sucrose at pH 7.4.*** *Significant transport of the drug through the PGPC vesicles can be detected in all experiments, visible by the gap in ΔI between the two time points.*

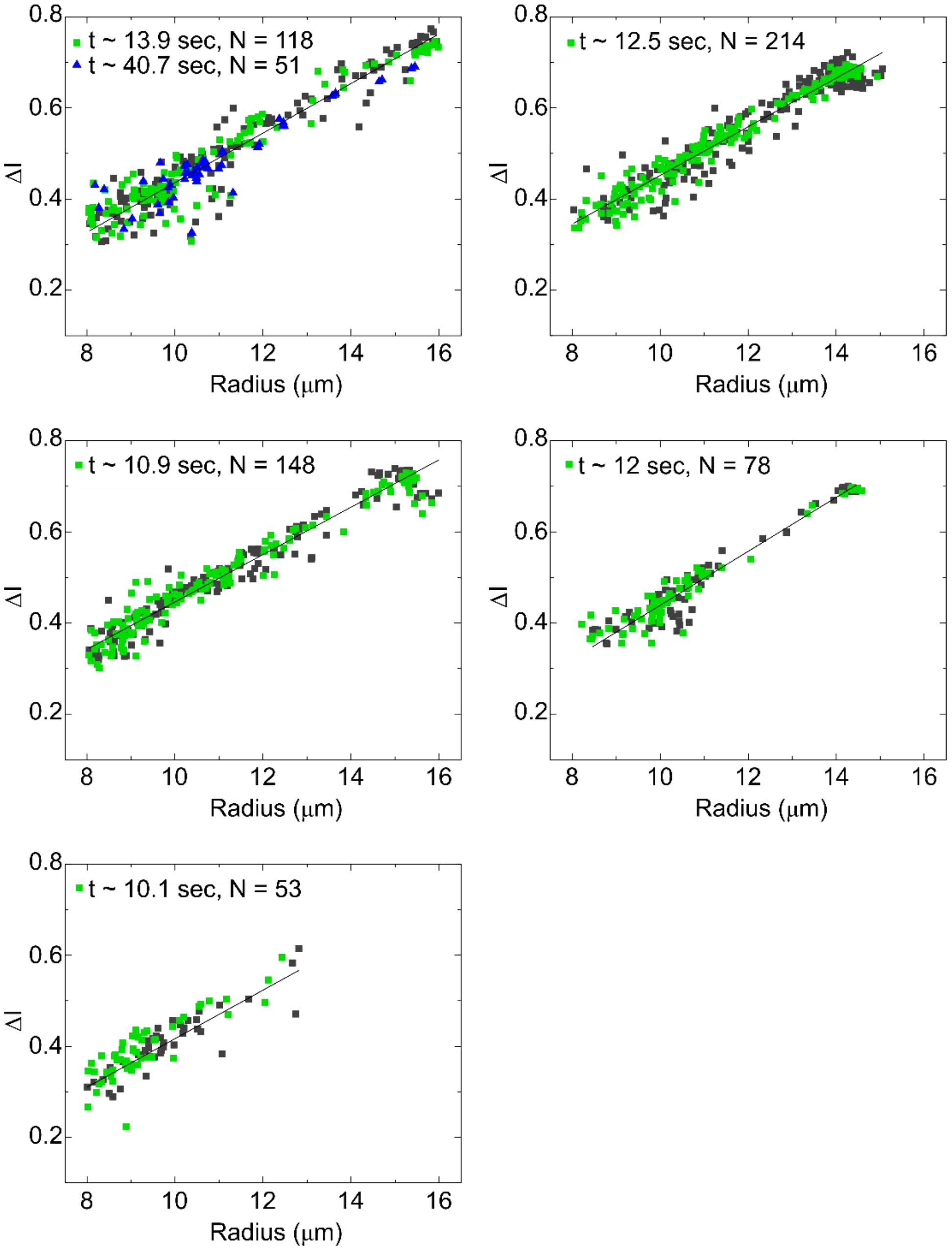


***Figure S4. Scatter plots of ΔI vs R for norfloxacin in 5 mM acetic acid buffer with 200 mM sucrose at pH 5.*** *No significant transport of the drug through the PGPC vesicles can be detected in any of the experiments, since no significant change of ΔI was detected between the different time points.*
